# COVID-19 in Space: Possible Health Risks and Preparedness Guidelines

**DOI:** 10.1101/2025.04.23.650113

**Authors:** Ishan Vashishat, Eddie Han, Barnabe D. Assogba

## Abstract

Background

The COVID-19 pandemic of 2020 resulted in over 705 million infections and more than 7 million deaths worldwide. The virus primarily spreads through aerosol droplets released during breathing, coughing, or sneezing, leading to symptoms ranging from mild fever and cough to severe outcomes, including death. Given the high risk associated with COVID-19, understanding its behaviour in diverse geographical and environmental conditions is critical. Space exploration and tourism represent an emerging industry, projected to reach a market value of $1.8 trillion. With numerous space missions planned by space agencies such as NASA, SpaceX, and ISRO, it is vital to address potential health risks for astronauts and space tourists.

**Objective:** With the expansion of human exploration into space, there is an urgent need to assess the risks posed by COVID-19 in extraterrestrial environments. This study reviews existing literature on airborne infections in space, identifies key knowledge gaps, and enhances preparedness for potential COVID-19 outbreaks during space missions.

**Methods:** A systematic literature review was conducted to identify studies examining airborne infectious diseases in space and their health effects under microgravity. Databases searched included PubMed and NASA’s Open Data Portal. To compare these findings with Earth-based data, additional systematic reviews were performed to analyze the known effects of these diseases on Earth, using Pathogen Safety Data Sheets. A separate systematic review was conducted using PubMed to explore similarities between COVID-19 and the selected airborne infectious diseases. Using a comparative approach, disease effects observed on Earth and in space were analyzed to predict COVID-19’s potential behavior in microgravity. Existing guidelines for managing airborne diseases in space and on Earth were reviewed and compared to develop a set of preparedness recommendations for COVID-19 in space.

**Results:** The airborne infectious diseases occurring in space found in this study include *Aspergillus fumigatus, Beauveria bassiana*, Epstein-Barr Virus (EBV), *Escherichia coli, Klebsiella pneumoniae* infections*, Pseudomonas aeruginosa*, Roseolovirus (Human Herpesvirus 6 & 7), *Salmonella Typhimurium* infection*, Serratia marcescens* infection*, Staphylococcus aureus, Staphylococcus epidermidis*, and Varicella-Zoster Virus (VZV). The relationship between the aforementioned diseases and COVID-19 was used in regard to theorizing the effects of COVID-19 in space. Six Tentative effects of COVID-19 in a microgravity environment could be theorized in this study. Along with that, recommendations to improve the current space travel health guidelines have also been referred to.

**Conclusion:** The results of this study will change the course of human space exploration by assisting in the protection of space travelers and guiding the development of new designs for spacecraft that include extra safety features.

## Introduction

Human space travel, which began in 1961, has seen 570 people journey beyond Earth’s atmosphere and return by 2021, with some individuals undertaking as many as seven missions during their careers [10]. The number of space travelers continues to grow, with 25 individuals living and working on the International Space Station in 2024, as reported by NASA [49]. However, a significant decline in space travel occurred in 2020 and 2021, with only 12 and 7 individuals traveling to space during those years, respectively, due to the disruptions caused by COVID-19 [10]. The COVID-19 pandemic led to global lockdowns and significant loss of human lives, severely affecting space operations and healthcare systems on Earth. If a similar outbreak were to occur during a space mission, it could jeopardize the mission’s success. The effects of COVID-19 in the extreme environment of microgravity remain largely unknown. Investigating these effects is crucial for developing guidelines to prevent future outbreaks and ensure the safety and success of space missions.

### COVID-19 as a viral vector

The year 2020 marks the emergence of COVID-19, an airborne infectious disease that triggered a global pandemic. Rapidly spreading across more than 200 countries, COVID-19 has infected over 776 million people worldwide, causing widespread social, economic, and healthcare disruptions [8, 51]. The name “COVID-19” which stands for Coronavirus Disease 2019, reflects its discovery in 2019. The disease is caused by SARS-CoV-2, a single-stranded RNA virus [8, 54]. COVID-19 is transmitted primarily through respiratory droplets and aerosols released when an infected person coughs, sneezes, or speaks [8]. Once inside the human body, the virus can manifest a wide range of symptoms. These can vary from mild, such as fever and cough, to severe, involving respiratory failure, septic shock, or even death [38].

As a viral infection, COVID-19 can disrupt the immune system, leading to various complications. One significant effect is heightened vulnerability to common secondary airway infections, often stemming from sleep disturbances caused by COVID-19 [48]. Additionally, COVID-19 infection can lead to a significant reduction in the number and function of natural killer (NK) cells, dendritic cells (DCs), and other immune cells [17]. This weakened immune response reduces the body’s ability to defend against other infections [47]. As a respiratory illness, COVID-19 disrupts oxygen utilization in the body, often leading to hypoxia. Microscopic analysis of lung tissue biopsies from COVID-19 patients reveals alveolar damage, lymphocytic inflammation, and edema [4]. Approximately 7.1% of affected individuals require mechanical ventilation due to these complication [26]. However, long-term use of ventilators is not without risks. Factors such as sedation, neuromuscular blockade, and over-ventilation can cause respiratory muscles to lose their workload, shifting it entirely to the ventilator. Over time, this may result in diaphragm thinning and atrophy [4]. Given COVID-19’s primary attack on the respiratory system, individuals with pre-existing respiratory conditions often experience heightened symptoms. For example, patients with chronic obstructive pulmonary disease (COPD) are at greater risk for severe complications, including thromboembolic events [45]. COVID-19 also worsens hypoxia in patients with underlying respiratory illnesses, intensifying symptoms and increasing the risk of severe outcomes, including death [45]. It has been observed that cold, dry conditions during winter can suppress immune function, thereby amplifying the effects of COVID-19 [28]. Similar environmental conditions are found in high-altitude regions and aboard space stations. Additionally, reduced exposure to sunlight and consequently lower levels of vitamin D can increase susceptibility to COVID-19 [6]. Such conditions are prevalent in dark environments, extreme valleys, and enclosed spacecraft.

### Incidence and Prevalence of COVID-19

In just one year, COVID-19 spread across the globe at an alarming rate. The officially reported death toll reached 1.8 million in 2020, though estimates suggest that the actual number of deaths may have exceeded 3 million worldwide [52]. Mutations in the virus continue to pose a significant global health threat. The latest strains currently in circulation as of April 14, 2025, are SARS-CoV-2 24C, 24F, and 24B [53]. COVID-19 has been shown to damage the olfactory epithelium, among other tissues, and its complex pathology makes it difficult to develop a comprehensive treatment, raising concerns about its eradication [14]. While acute COVID-19 symptoms typically last up to four weeks, cases of long-term COVID-19 are increasingly reported. Long-term COVID, also known as “post-acute sequelae of COVID-19” (PASC), refers to the persistence of one or more COVID-related symptoms for an extended period after recovery [13]. About 57% of patients report at least one lingering symptom even a year after treatment [33]. The most commonly reported symptoms include dyspnea on exertion (34%), difficulty concentrating (32%), fatigue (31%), frailty (31%), and arthromyalgia (28%) [33]. Even after recovering from COVID-19, individuals can develop multiple health complications. These range across various systems, including the pulmonary, cardiovascular, neurological, hematological, inflammatory, renal, endocrine, gastrointestinal, and integumentary systems [7]. Common symptoms among long COVID patients include persistent fatigue, pneumonia, cardiac dysfunction, nonrestorative sleep, and multi-inflammatory syndrome (MIS) [7]. Due to the relatively short time since the initial COVID-19 outbreak, research is still ongoing, and not all long-term effects have been fully identified. Furthermore, the incidence and prevalence of COVID-19 in space or microgravity conditions remain unknown. However, diseases such as Epstein-Barr virus have demonstrated increased virulence and prolonged survival under microgravity conditions [31].

Geographical factors have significantly influenced the spread and severity of COVID-19. For instance, the average time from symptom onset to hospitalization was 5.7 days in mainland China, compared to 3.3 days outside of China [26]. This discrepancy suggests that the incubation period may vary depending on the location where the virus is contracted. Climate has also played a key role in the spread of the virus. New strains of COVID-19 have been observed to emerge between June and September in the southern hemisphere, in countries like Argentina, Australia, Brazil, and New Zealand [1]. Furthermore, weather conditions greatly influence transmission rates. Cold and dry environments are associated with higher transmission rates, whereas warm and humid conditions tend to slow the spread of the virus [30]. While any single factor might not significantly affect the spread of COVID-19, the combination of multiple meteorological factors has a pronounced impact. Models incorporating several weather-related variables have been shown to more accurately describe epidemic trends compared to single-factor models [9].

### Space and Space Travel

Given the variation in COVID transmission and symptoms across the world, there are serious questions regarding disease progression in extreme environments. One of the most extreme environments encountered by humans is space. Space is defined as the area above 100 km above the earth’s surface [20]. Space stations and space shuttles are considered extreme environments because of their microgravity. Space travel has seen significant growth since its inception and is projected to expand further in the future. As of 2021, 570 individuals had travelled to space and returned [10]. The peak year for human space travellers was 1985, when 63 individuals ventured into space. However, the COVID-19 pandemic led to a sharp decline, with only 9, 12, and 7 individuals travelling to space in 2019, 2020, and 2021, respectively [11]. Although it may seem like space travel has decreased in recent years, the trend is now shifting. In 2024 alone, NASA reported that 25 people lived and worked in space, highlighting the growing demand for space travel in the post-pandemic era.

The demand for and necessity of space exploration have surged as technological advancements continue. In recent years, both private and public interest in space travel has grown dramatically. Companies like SpaceX, Virgin Galactic, and Blue Origin have been catering to an emerging market, with the potential to surpass the aviation industry in terms of market share [43]. In June 2024, SpaceX made headlines by launching four civilians on a space tour, marking the dawn of civilian space travel [35]. By 2035, space economy is expected to be worth $1.8 trillion, due to increasing prevalence of satellite and space technology [23]. In addition, multiple countries have announced future space missions and projects. For example, China has outlined an ambitious 20-year plan to establish lunar factories, laboratories, and tourism facilities. This initiative includes a 10-person crew landing with provisions for 100% *in-situ* oxygen and water production [21]. Similarly, India is making significant strides in space exploration. Under its Gaganyaan program, India plans to launch eight manned missions starting 2028 and initiate the development of a space station in the same year [2].

Additionally, space mining has gained interest as a viable future option for harvesting resources. Private firms like SpaceX and Blue Origin are reducing launch costs and developing more sustainable spacecraft capable of performing multiple flight cycles [5]. These advancements make mining in space increasingly feasible. The potential benefits are immense—space mining could help mitigate climate change on Earth and prevent the depletion of natural resources. Metallic asteroids frequently hold significantly higher concentrations of minerals compared to Earth. For instance, iron, cobalt, and nickel concentrations on asteroids are on avergae 893,000 g/mt, 6,000 g/mt, and 93,000 g/mt, respectively, compared to only 41,000 g/mt, 20 g/mt, and 80 g/mt in an open pit mine on Earth [12]. These untapped resources represent a promising incentive for space exploration.

### Effects of Space Travel on the Human Body

Space travel presents unique challenges to human health. From a psychological standpoint, isolation and confinement during space missions can take a significant toll. The small, enclosed environment of a spacecraft can also become claustrophobic, potentially leading to interpersonal conflicts [27]. Moreover, without Earth’s atmospheric protection, astronauts are exposed to high levels of space radiation. This radiation comes from the Earth’s magnetic field, solar particles, and galactic cosmic rays, increasing the risk of cancer and degenerative diseases such as heart disease and cataracts [32]. In the absence of gravity, astronauts experience muscle atrophy, with muscle mass deteriorating over time. Bone health is similarly impacted, as astronauts lose between 1% to 1.5% of bone mineral density per month in space [44]. This bone loss further increases the complications of long-term flights. When calcium is drained from bones into the rest of the body, the kidneys have to bear the extra burden of excreting the calcium, increasing the risk of astronauts developing kidney stones [46].

The effects of space travel extend to the immune system. Immune cells generated in space show altered gene expression. B cells, CD4+ T cells, CD8+ T cells, and CD14+ monocytes are all influenced by Differentially Expressed Genes (DEG) [22]. Additionally, T cells (both CD4+ and CD8+) exhibit heightened responsiveness to an infection, while B cells, essential for antibody production, become less responsive [35]. A more detailed study of immune functions in space is currently limited because the effects of these DEGs return to normal shortly after astronauts return to Earth [22]. The microgravity environment of space can surprisingly benefit lung function. Cardiac output to the lungs increases by approximately 28%, enhancing the absorption of gases [37]. However, respiratory diseases remain a challenge during space missions. One notable respiratory disease outbreak occurred during the Apollo 7 mission, when three astronauts contracted a respiratory infection that spread rapidly and disrupted mission cooperation [36]. The infection originated with astronaut Wally Schirra and quickly spread to the crew. The biggest complication was the accumulation of phlegm, which had to be suctioned out due to the absence of gravity [34]. Preventive measures, such as increasing absolute humidity (AH) inside spacecraft, may help in reducing viral transmission. This approach is supported by findings from Earth, where higher AH levels are linked to decreased viral spread [50].

### Knowledge Gap

There is a significant lack of research exploring the specific risks COVID-19 poses in space environments. This gap encompasses both the potential continuation of COVID-19 infections contracted on Earth and the possibility of new infections occurring in space due to the unique environmental conditions.

Even though more than 570 individuals have traveled to space, our understanding of how diseases and viruses behave in microgravity and isolated environments remains limited. The COVID-19 pandemic underscores the urgency of addressing these unknowns. Understanding the behavior of infectious diseases like COVID-19 in space is critical for increased space exploration and the numbers of humans traveling to space is likely to increase dramatically in the near future. Comprehensive knowledge in this area could help mitigate risks, ensure mission success, and prevent unnecessary fatalities during long-duration space travel.

### Research Questions

The present study seeks to explore two key questions. First, what are the possible increased risks COVID-19 poses to human health during space travel? Second, what preparedness measures can be implemented to prevent COVID-19 outbreaks in space environments? Addressing these questions will provide valuable insights and contribute to the development of disease management protocols essential for safeguarding the health of astronauts and ensuring the success of future space missions.

To understand the potential risks of COVID-19 symptoms in space, it is important to explore its similarities with other airborne infectious diseases. These similarities could include the involvement of immune responses, such as T cells and B cells, physiological reactions, and changes in cell counts. Investigating how airborne infectious diseases behave in space environments is also essential, including identifying which infectious diseases have previously affected astronauts. By comparing the impact and viability of various diseases in space, we can estimate the potential risk posed by COVID-19. For instance, if another disease exhibits similar effects on both Earth and in space, it is reasonable to predict that COVID-19 may behave similarly as well, though the intensity of its symptoms might differ. On the other hand, if certain symptoms appear only on Earth and not in space, we can speculate that COVID-19 might follow a similar pattern.

To minimize the risk of COVID-19 transmission in space, developing a preparedness plan is crucial. This can involve reviewing existing guidelines for managing airborne diseases in space and determining whether these protocols align with those used on Earth. If they do, similar measures could be adapted to address the transmission of COVID-19 during space missions. Drawing on previously established guidelines will provide a structured framework for preventing the spread of the virus in the unique conditions of space.

This study aims to address the lack of research on the behavior of SARS-CoV-2 in space by examining other airborne infectious diseases previously studied in microgravity. Through systematic literature reviews, our research will identify patterns and similarities between airborne infectious diseases studies in space and COVID-19 to predict potential risks to astronauts. Comparisons will be drawn between the effects of previously identifies airborne diseases in space and on Earth to hypothesize how COVID-19 might behave in a microgravity environment. Our findings will help in the development of a comprehensive preparedness plan for COVID-19 prevention through integrating existing guidelines for prevention of airborne infections in space and on Earth, alongside Earth-based COVID-19 protocols, to create evidence-based measures for managing COVID-19 during space missions.

## Methods

### Systematic Review for studies of Airborne Infectious Diseases in Space

A systematic search of the literature was conducted to identify articles investigating airborne infectious diseases in space and their effects in a microgravity environment. The databases searched included PubMed, NASA’s Open Data Portal, and ScienceDirect. However, due to time constraints, ScienceDirect was not used in the final review. No restrictions were placed on the publication date of the articles. The search strategy included the following keywords: (“Airborne Diseases” OR “Airborne Infection” OR “Airborne” OR “Infectious” OR “Infection”) AND (“Spaceflight” OR “Microgravity” OR “Space Environment”).

#### Inclusion and Exclusion Criteria

Studies were included if they were conducted in space, featured an airborne infectious disease or pathogen, and described the health effects of the disease in a space environment. Due to the limited availability of relevant studies, generalized effects of the identified diseases and pathogens were also considered. Secondary factors, such as whether the study was a primary research article or whether it presented future safety guidelines, were also noted during the inclusion process; however, these factors were not prerequisites for inclusion. Articles were excluded if they were inaccessible or written in a language that posed a barrier to comprehension, though no open-source or language filters were applied in the initial searches. The selection process was facilitated using an assessment form.

#### Data Extraction

Following the assessment, relevant and accessible articles were used for data extraction. The extracted data included information on the identified diseases, the reported health effects, the number of recorded cases (if available), and the guidelines implemented for managing these diseases in space. Initially, data were collected using an extraction form (Appendix) and later organized into an Excel spreadsheet, where studies were categorized based on the respective diseases examined.

#### Collection the effects of Airborne Infectious Diseases on Earth

To ensure a fair comparison, the health effects of these diseases in space were compared to their known effects on Earth. For standardization, reference data were obtained from the Pathogen Safety Data Sheets: Infectious Substances published by the Government of Canada. The absence of standardized Pathogen Safety Data Sheets resulted in the exclusion for three out of twelve diseases (Figure 1).

**Figure 1:**
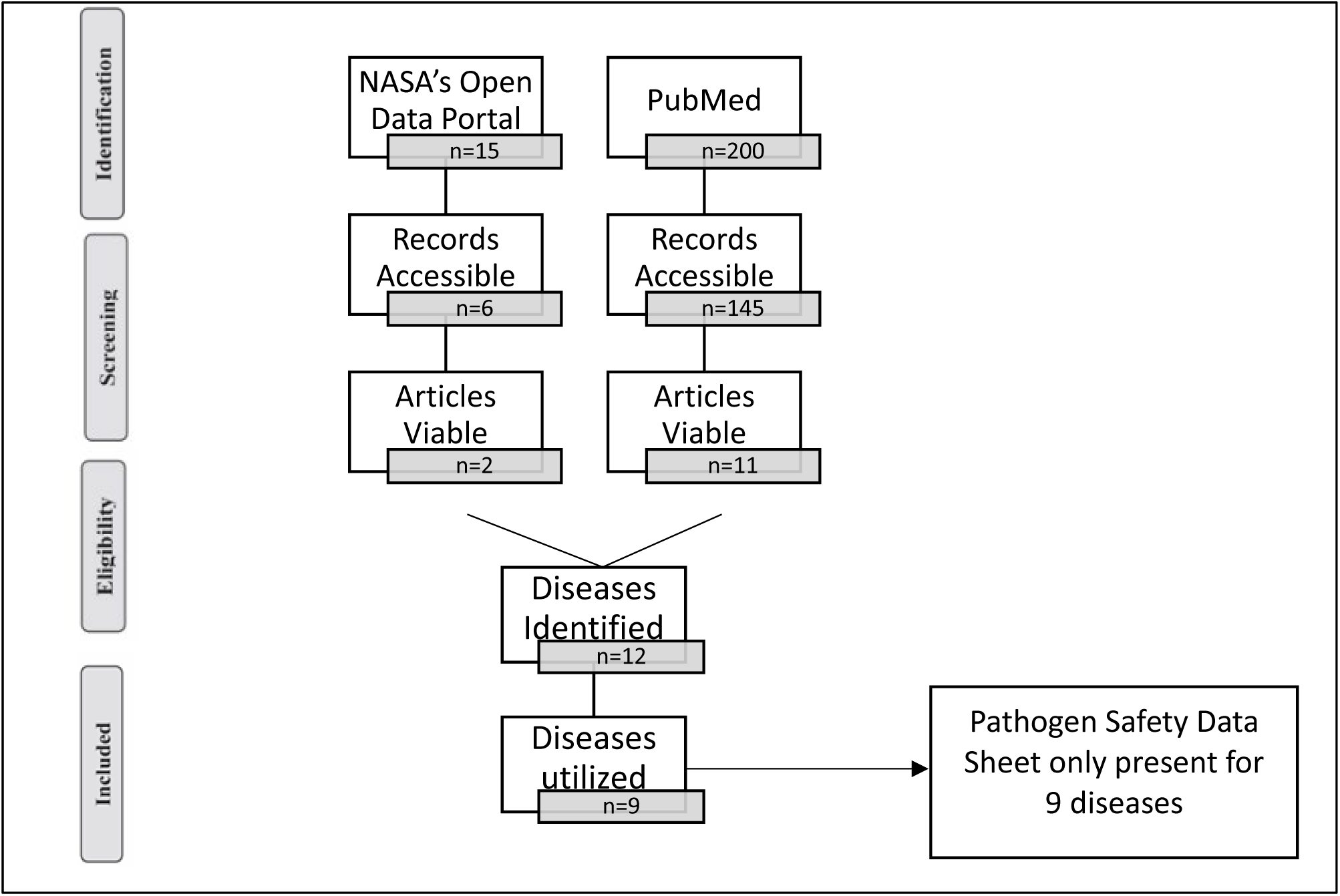
Flowchart depicting the systematic review process for identifying, screening, and selecting studies on airborne infectious diseases in space. The diagram illustrates the number of records retrieved from NASA’s Open Data Portal and PubMed, the number of accessible records, viable articles, identified diseases, and the final number of diseases included in the study. The exclusion of three diseases was due to the absence of standardized Pathogen Safety Data Sheets.

### Systematic Review for studies that link Airborne Infectious Diseases to COVID-19 on Earth

A systematic search of the literature was conducted to identify articles reporting similarities between COVID-19 and other airborne infectious diseases previously identified as they manifest on Earth. This data collection resulted in a series of nine systematic reviews (Figure 1). The database used for this phase of the systematic review was PubMed. A time restriction was applied, and any article published before 2020 was excluded. The search strategy included the following keywords: (“COVID-19” OR “Coronavirus” OR “COVID”) AND *-disease-*, where *-disease-* was systematically replaced with one of the following: *Salmonella typhimurium*, *Serratia marcescens*, Varicella-Zoster Virus, *Aspergillus fumigatus*, Epstein-Barr Virus, *Pseudomonas aeruginosa*, *Escherichia coli*, *Staphylococcus aureus*, and *Klebsiella pneumoniae*—the diseases identified in the initial search. The search results were exported into an Excel spreadsheet, and each accessible article was stored as a PDF file for future reference. Titles and abstracts were screened using the inclusion criteria listed below, and the full texts of potentially relevant articles were obtained for further review.

#### Inclusion and Exclusion Criteria

Studies were included if they discussed COVID-19, addressed the secondary disease, and mentioned how their symptoms coincide. Secondary factors, such as whether the study was a primary research article, were also noted during the inclusion process; however, these factors were not prerequisites for inclusion. Articles were excluded if they were inaccessible or written in a language that posed a barrier to comprehension, though no open-source or language filters were applied in the initial searches. The selection process was facilitated using an assessment form (Appendix).

#### Data Extraction

Following the assessment, relevant and accessible articles were used for data extraction. The extracted data included information on the similarities between COVID-19 and the secondary disease. Initially, data were collected using an extraction form (Appendix 4) and later organized into an Excel spreadsheet, where studies were categorized based on the respective diseases examined.

### Estimate the Risk for COVID-19 in Space

To predict the potential effects of COVID-19 in space, a comparative approach was employed, utilizing data from airborne infectious diseases that have been studied in both terrestrial and microgravity environments. If an infectious disease exhibited specific health effects on both Earth and in space, this provided a foundation for hypothesizing how COVID-19 might behave in a space environment.

The information from all three parts of the study: disease characteristics in space, disease characteristics on Earth, and COVID-19’s comparative effects was analyzed using data compiled in an Excel spreadsheet. Specifically, the analysis was if COVID-19 shared similar physiological impacts with a known disease on Earth, and the effects of that disease had been studied in space, it allowed for a reasonable extrapolation of COVID-19’s potential pathophysiology, transmission, and risks under microgravity conditions. The comparison of these datasets enabled the theorization of speculated health effects and generalized physiological impacts of COVID-19 in a microgravity environment. This approach facilitated a structured hypothesis regarding COVID-19’s possible behavior, transmission, and impact on astronaut health in space, despite the absence of direct experimental data on the virus in spaceflight conditions.

### Develop Preparedness Guidelines against COVID-19 in Space

The current guidelines for managing airborne infectious diseases in space were first reviewed. This was done using the same set of studies identified in the earlier phases of the research to examine established protocols. These protocols were compiled and organized into a tabulated format to create a structured reference for existing disease control measures in microgravity environments.

To complement this, the guidelines for managing the same airborne infectious diseases on Earth were systematically gathered and stored in a database. Reference data were obtained from the Pathogen Safety Data Sheets: Infectious Substances published by the Government of Canada along with Canadian Biosafety Handbook, providing a standardized source of terrestrial disease management protocols. Following this, existing guidelines for managing COVID-19 on Earth were also incorporated into the database, ensuring a direct comparison with protocols used for other airborne diseases. Specifically, the Pathogen Safety Data Sheets: Infectious Substances for COVID-19, published by the Government of Canada, was utilized as a key reference.

The final step involved a comparative analysis of three datasets: disease management protocols in space, terrestrial guidelines for other airborne infectious diseases, and COVID-19-specific guidelines on Earth. To ensure consistency in guideline identification, all pathogens selected for comparison were classified according to their Risk Group and Containment Level using the Public Health Agency of Canada’s Pathogen Safety Data Sheets. Guidelines corresponding to each classification were sourced from the *Canadian Biosafety Handbook*. These classifications were used to standardize the containment and response measures evaluated later in the study. For the purpose of comparison, COVID-19 was also categorized based on its listing in the Pathogen Safety Data Sheets, which places it under Risk Group 2 and Containment Level 2. By identifying patterns and similarities across these three sources, a set of potential guidelines for preventing the spread of COVID-19 in space was proposed.

## Results

### Systematic Review for Studies of Airborne Infectious Diseases in Space

A systematic review of the literature was conducted to identify articles investigating airborne infectious diseases in space and their effects in a microgravity environment. The databases searched included PubMed and NASA’s Open Data Portal. The search strategy included the following keywords: (“Airborne Diseases” OR “Airborne Infection” OR “Airborne” OR “Infectious” OR “Infection”) AND (“Spaceflight” OR “Microgravity” OR “Space Environment”). No restrictions were placed on the publication date of the articles. The systematic review yielded 215 studies, the titles and abstracts of which were screened for the inclusion criteria. Searches were conducted using databases such as PubMed and NASA’s Open Data Portal. Figure 1 illustrates the study selection process via a flow diagram. Of the total results, 30% (64 studies) were excluded due to inaccessibility, either because they lacked a digitally accessible version, were restricted by language, or were locked behind paywalls. The remaining 151 studies were assessed for eligibility as per the inclusion criteria, from which 13 studies were deemed viable for further data extraction.

From these 13 studies, 12 airborne infectious diseases were identified, and their effects in space environments were extracted and are summarized in Table 2. However, only 9 out of the 12 diseases had corresponding Pathogen Safety Data Sheets (PSDS) published by the Government of Canada, which were used to compare the effects of the diseases on Earth. As a result, 3 diseases were excluded from the current analysis due to the absence of standardized terrestrial comparison data. The effects of the remaining 9 diseases, both in space and on Earth — are detailed in Table 2.

### Systematic Review for studies that link Airborne Infectious Diseases to COVID-19 on Earth

To compare the effects of selected airborne infectious diseases with those of COVID-19 on Earth, a series of systematic reviews were conducted. PubMed was used as the primary database for this phase of the review. A time restriction was applied to exclude articles published before 2020, in order to better align the findings with the post-pandemic context of COVID-19. The search strategy included the following keywords: (“COVID-19” OR “Coronavirus” OR “COVID”) AND *-disease-*, where *-disease-* was systematically replaced with one of the following: *Salmonella typhimurium*, *Serratia marcescens*, *Varicella-Zoster Virus*, *Aspergillus fumigatus*, *Epstein-Barr Virus*, *Pseudomonas aeruginosa*, *Escherichia coli*, *Staphylococcus aureus*, and *Klebsiella pneumoniae*, the diseases identified in the initial search. These searches yielded a total of 4,124 studies (Figure 2).

**Figure 2:**
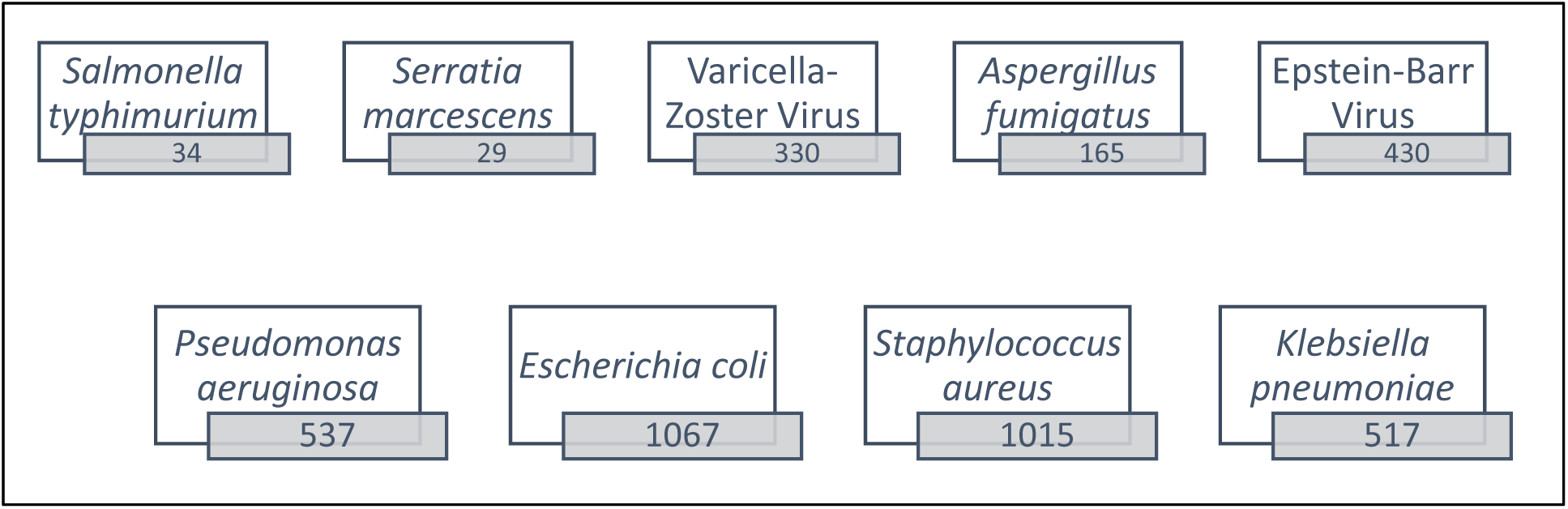
**Number of articles retrieved from PubMed for each airborne infectious disease in comparison to COVID-19**. The search strategy included the combination of COVID-19 with each listed disease, resulting in varying numbers of relevant articles. *Escherichia coli* and *Staphylococcus aureus* had the highest number of retrieved studies, while *Serratia marcescens* had the lowest.

The diseases included in the comparison were *Salmonella typhimurium, Serratia marcescens*, and Epstein-Barr Virus (EBV). For *Salmonella typhimurium*, 34 studies were identified, of which 25 were accessible. However, none were found to be in line with the inclusion criteria, either missing COVID-19, *Salmonella typhimurium,* or a link between them, for extracting comparative data with COVID-19. For *Serratia marcescens*, 29 studies were identified, with 27 accessible. Among these, 1 study was deemed viable, showing some degree of correlation between *Serratia marcescens* and COVID-19. For Epstein-Barr Virus, 430 studies were identified, 401 accessible. Among those 21 were deemed viable, showing some degree of correlation between Epstein-Barr Virus and COVID-19. This ongoing review is illustrated in Figure 3. The comparative findings for the selected diseases and COVID-19 are summarized in Table 2.

**Figure 3:**
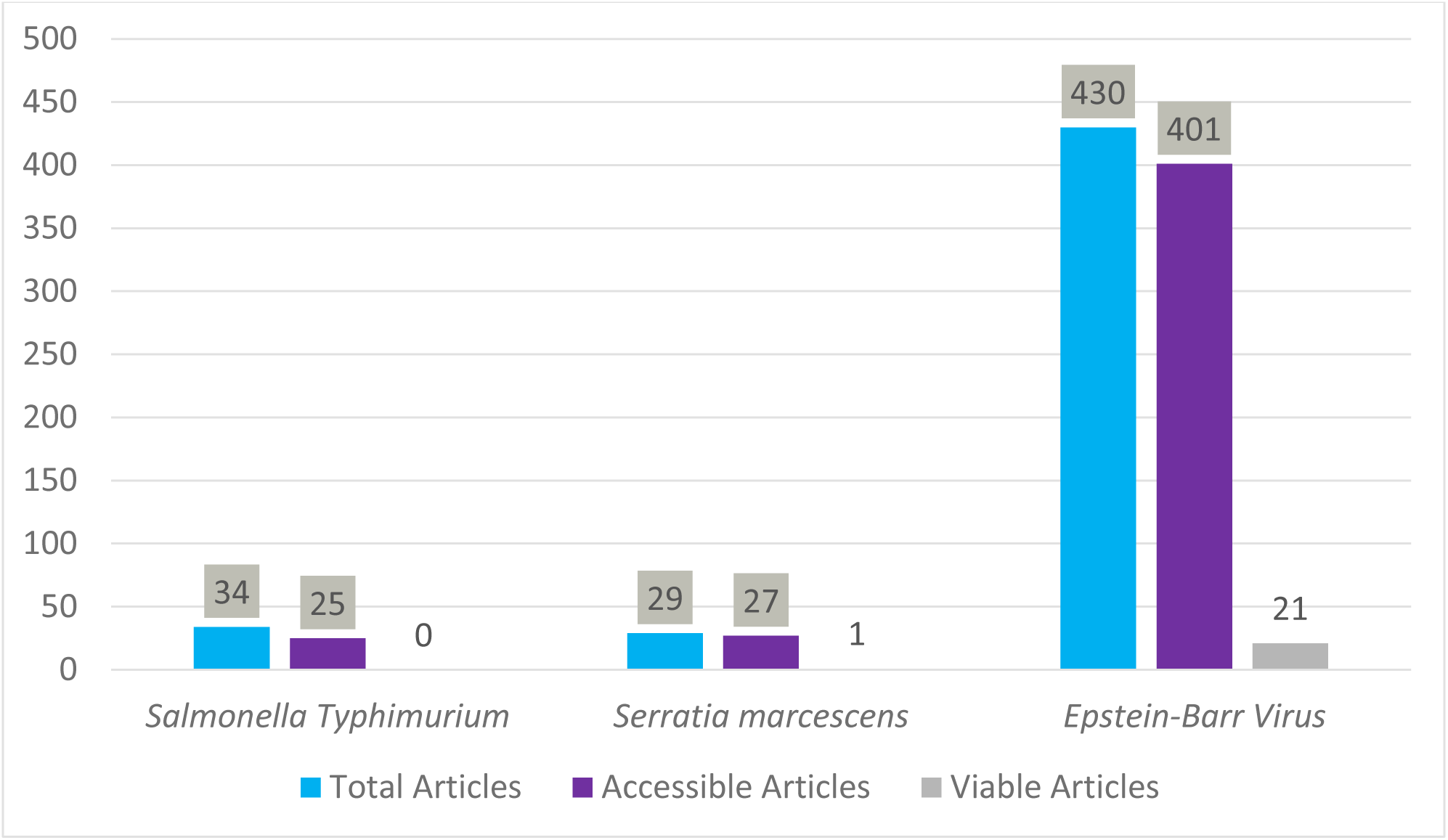
Literature availability on possible studies comparing COVID-19 to different pathogens. Number of studies from the systematic review used to establish a link between COVID-19 and the airborne infectious diseases identified earlier. The figure compares the total, accessible, and viable research articles on Salmonella typhimurium, Serratia marcescens, and Epstein-Barr Virus. “Viable articles” refer to those that meet the inclusion criteria.

### Estimate the Risk for COVID-19 in Space

Using comparative analysis, if COVID-19 was found to share similar physiological impacts with a known disease on Earth—one that had already been studied in space—it allowed for a reasonable extrapolation of COVID-19’s potential pathophysiology, transmission, and associated risks in microgravity. This comparison of datasets enabled the formulation of theoretical health effects and generalized physiological outcomes of COVID-19 in a space environment. By comparing the effects of COVID-19 to those of Epstein-Barr Virus and Serratia marcescens, six potential effects of COVID-19 in space were identified: Immune Suppression & Reactivation, Inflammatory Response, Neurocognitive and Systemic Symptoms, Gastrointestinal and Pulmonary Risk, Increased Virulence & Growth Potential, and Greater Severity & Mortality Risk (Table 1).

**Table 1:** Predicted effects of COVID-19 in space based on comparisons with Epstein-Barr Virus and *Serratia marcescens*. This table outlines six potential health impacts of COVID-19 in space, derived from observed similarities with the behavior of EBV and Serratia marcescens under spaceflight or microgravity conditions. These effects include immune suppression and reactivation, heightened inflammatory responses, neurocognitive and systemic symptoms, gastrointestinal and pulmonary risk, increased virulence and growth potential, and greater severity and mortality risk.

|  |  |
| --- | --- |
| **Immune Suppression & Reactivation:** | Like EBV, COVID-19 may exhibit increased reactivation or persistence in space due to spaceflight-induced immune suppression, leading to prolonged viral activity and increased risk of complications such as inflammation or post-viral syndromes. |
| **Inflammatory Response:** | As seen with both EBV and *Serratia marcescens*, COVID-19 may trigger heightened inflammatory responses in space, potentially worsening outcomes such as fever, fatigue, or organ involvement (e.g., lungs, liver). |
| **Neurocognitive and Systemic Symptoms:** | Based on similarities with EBV's association with fatigue, brain fog, myalgia, and cognitive dysfunction, COVID-19 in space may similarly lead to long COVID-like symptoms affecting neurological and muscular systems. |
| **Gastrointestinal and Pulmonary Risk:** | Like EBV's role in gastrointestinal inflammation and *Serratia*'s pulmonary risks, COVID-19 may also result in exacerbated GI and respiratory issues in space. |
| **Increased Virulence & Growth Potential:** | As *Serratia marcescens* demonstrated enhanced virulence due to altered bacterial growth kinetics in microgravity, COVID-19's replication rate and tissue tropism could be altered, potentially amplifying disease severity. |
| **Greater Severity & Mortality Risk:** | Given that both EBV and *Serratia* infections are associated with worse clinical outcomes in space or co-infection settings, COVID-19 in space may lead to greater severity, especially in critically ill or immune-compromised astronauts. |

### Develop Preparedness Guidelines against COVID-19 in Space

Current guidelines for managing airborne infectious diseases in space were extracted from the same 13 viable studies referenced earlier (Figure 2). These studies provided insight into the recommended procedures and safety measures implemented in space environments when dealing with these infectious agents. In order to establish a common ground, each identified pathogen was cross-referenced with its corresponding Pathogen Safety Data Sheet provided by Government of Canada [40]. According to the data obtained, all of the pathogens were classified as Risk Group 2 and required Containment Level 2 precautions. The containment guidelines associated with Risk Group 2 and Containment Level 2 were sourced from the Canadian Biosafety Handbook. Since all the pathogens identified in the space studies fell within this classification, it was assumed that the operational guidelines followed for these diseases in space mirror those recommended for Containment Level 2 organisms. COVID-19, according to Pathogen Safety Data Sheet, also falls under Risk Group 2 and Containment Level 2 (Table 3). Therefore, based on risk equivalence, it was inferred that the guidelines for managing COVID-19 in space should closely align with those already in place for other airborne infectious diseases identified in this review.

**Table 2:** Comparison of health effects of airborne infectious diseases in space and on Earth, and their relevance to COVID-19. This table summarizes the known effects of 12 airborne pathogens in both space and Earth environments, including *Salmonella typhimurium, Serratia marcescens, Varicella-Zoster Virus, Aspergillus fumigatus, Epstein-Barr Virus, Pseudomonas aeruginosa, Escherichia coli, Staphylococcus aureus, and Klebsiella pneumoniae*. It highlights similarities in virulence, immune response, and symptom overlap with COVID-19, where applicable. The table also notes where reviews were not conducted due to time constraints or limited data availability.

| Sr. No. | Disease | Effects in Space | Effects on Earth | Comparison to COVID-19 |
| --- | --- | --- | --- | --- |
| 1 | Salmonella Typhimurium infection | Increased virulence and stress resistance of Salmonella in spaceflight conditions.<br>Altered immune and gastrointestinal responses in human intestinal epithelial cells<br>Potential for heightened infection risk during space missions.<br>Space-grown strains showed hypervirulence in murine models.<br>Increased biofilm formation under low-shear microgravity conditions.<br>S. Typhimurium exhibited increased virulence and stress resistance in spaceflight conditions<br>Enhanced pathogen colonization and altered transcriptional responses in infected human cells<br>Spaceflight altered host immune responses, including reduced TLR4 expression, which may impair immune recognition of Salmonella<br>Infection triggered higher pro-inflammatory responses in | Salmonella Typhimurium commonly causes gastroenteritis, which begins suddenly with nausea, vomiting, abdominal cramps, diarrhea, headache, chills, and fever up to 39°C.<br>The symptoms of gastroenteritis usually last between 5 to 7 days and can range from mild to severe.<br>It is responsible for approximately 1.3 billion cases and 3 million deaths globally each year, including 1.4 million cases and 600 deaths in the U.S. annually.<br>In about 3–10% of infected people, the infection progresses to bacteremia, where the bacteria enter the bloodstream.<br>Bacteremia can lead to septic shock and endocarditis, particularly in people over 50 or those with existing heart conditions.<br>It can also cause infections of the aorta, liver, spleen, and biliary tract, especially in those with structural abnormalities. | No Connection Established |
|  |  | spaceflight-exposed human cells compared to ground controls | Other complications include mesenteric lymphadenitis, osteomyelitis (especially in long bones and the spine), urinary tract infections, pneumonia, pulmonary or brain abscesses, subdural and epidural empyema, and meningitis (rarely).<br>In severe cases, death may occur as a result of bacteremia.<br>People with weakened immune systems or pre-existing conditions like HIV/AIDS, diabetes, cancer, cirrhosis, chronic granulomatous disease, or sickle cell disease are at higher risk of developing these serious complications. |  |
| 2 | *Serratia marcescens* infection | Increased virulence of *Serratia marcescens* under spaceflight and microgravity conditions, leading to decreased survival rates.<br>The increased lethality is linked to bacterial growth kinetics rather than changes in host immune response.<br>Increased virulence of *Serratia marcescens* when grown in spaceflight conditions, leading to decreased survival rates in *Drosophila melanogaster*.<br>The increased lethality was linked to changes in bacterial growth kinetics rather than host immune suppression. | *Serratia marcescens* is an opportunistic pathogen and one of the top ten causes of bacteremia in North America.<br>It can cause a wide range of infections including bacteremia, pneumonia, intravenous catheter-associated infections, osteomyelitis, endocarditis, and in rare cases, endophthalmitis (an eye infection).<br>Endophthalmitis symptoms appear rapidly and may include fever, redness, ocular pain, swelling around the eyes, and pus in the eye (hypopyon). | Infections can lead to bacterial cross-contamination<br>Increased risk of *S. marcescens* superinfection<br>Ventilator-associated pneumonia and septic shock. |
|  |  |  | The mortality rate due to Serratia bacteremia can reach 37% six months post-infection.<br>Other rare neonatal infections include otitis externa, enterocolitis, omphalitis, gastroenteritis, septic arthritis, and intraperitoneal abscesses.<br>Risk factors in neonates include low birth weight, use of mechanical ventilation, and premature birth (under 37 weeks).<br>The mortality rate in neonates with Serratia infection is reported to be as high as 44%.<br>Most infections are caused by non-pigmented strains, which are more virulent than pigmented ones. |  |
| 3 | Varicella-Zoster Virus (VZV) | Reactivation risk increases in spaceflight.<br>Higher viral load in astronauts, increasing transmission potential.<br>Causes chickenpox (more easily spread) and shingles<br>Asymptomatic VZV reactivation was detected in astronauts. | Causes a generalized itchy, vesicular rash and fever, primarily in children.<br>Lesions typically begin on the back of the head and ears, spreading to the face, neck, trunk, and limbs.<br>The number of lesions can range from 10 to several hundred.<br>May lead to viremia and visceral organ involvement in immunocompromised individuals. | Review not conducted due to time constraints. |
|  |  | Spaceflight stress suppressed cell-mediated immunity (CMI), leading to higher viral shedding.<br>VZV DNA was present in astronaut saliva during and after spaceflight, confirming reactivation without visible disease | Can cause pneumonia, hepatitis, encephalitis, transient ataxia, aseptic meningitis, and transverse myelitis.<br>Thirty percent of children with leukemia may develop disseminated varicella, with a 7% mortality rate.<br>Less common complications include arthritis, glomerulonephritis, myocarditis, and purpura fulminans.<br>The virus becomes latent in sensory ganglia neurons after initial infection.<br>Reactivation of the virus leads to shingles (herpes zoster), especially in people over 50 years old.<br>Shingles causes a painful vesicular rash along the area of skin supplied by the affected sensory nerve.<br>Postherpetic neuralgia may occur, with pain lasting months or years after the rash heals.<br>In immunocompromised patients, reactivation can lead to disseminated infection involving visceral organs. |  |
| 4 | Aspergillus fumigatus | Causes aspergillosis, which is a serious infection in immunocompromised individuals or those with chronic lung conditions. | Aspergillus fumigatus is the most common cause of aspergillosis, a group of diseases caused by Aspergillus spp. | Review not conducted due to time constraints. |
|  |  | Space conditions may enhance its virulence | It can cause allergic reactions such as allergic bronchopulmonary aspergillosis, rhinitis, and farmer's lung.<br>It may result in superficial and local infections like cutaneous infections, otomycosis (ear infection), and tracheobronchitis.<br>Aspergilloma (fungal ball in the lungs) and osteomyelitis (bone infection) can occur in patients with damaged tissue.<br>Invasive pulmonary aspergillosis is the most severe form, primarily affecting immunocompromised individuals.<br>Invasive infections often start in the sinopulmonary tract, particularly the lungs.<br>Invasive sinusitis symptoms include fever, facial pain, headache, swelling, nosebleeds, proptosis, nerve issues, and palate ischemia.<br>Invasive pulmonary aspergillosis presents with fever, cough, dyspnea, and may include pleural chest pain due to vascular invasion. |  |
|  |  |  | If left untreated, the infection can spread through the bloodstream to other organs.<br>Central nervous system involvement can lead to seizures or stroke.<br>The mortality rate in immunocompromised patients can be significant, especially without early treatment.<br>High-risk groups include patients with acute leukemia, transplant recipients, and those with neutropenia or graft-versus-host disease. |  |
| 5 | Beauveria bassiana | Can cause disseminated infections, though it is rarely a human pathogen | Pathogen Safety Data Sheet Not Available | Review not conducted due to time constraints. |
| 6 | Roseolovirus (Human Herpesvirus 6 & 7) | Known to cause fever, rash, and neurological complications.<br>Potential reactivation in astronauts due to spaceflight-induced immune suppression | Pathogen Safety Data Sheet Not Available | Review not conducted due to time constraints. |
| 7 | Epstein-Barr Virus (EBV) | Causes infectious mononucleosis (mono).<br>Increased viral shedding in saliva under spaceflight conditions<br>Space radiation exposure led to reactivation of latent EBV, increasing viral load in astronauts. | Most childhood EBV infections are asymptomatic.<br>EBV can cause infectious mononucleosis.<br>Symptoms of infectious mononucleosis include fever, sore throat, pharyngitis, tonsillitis, lymphadenopathy, and fatigue. | Participants with long COVID showed elevated antibody responses to EBV antigens, especially p23 and gp42, suggesting EBV reactivation. This reactivation is linked to symptoms such as fatigue and immune dysfunction, which are also seen in long COVID, indicating similar post-viral sequelae. |
|  |  | Lytic gene transcription was induced at all radiation doses, with the highest activation at day 4 post-exposure.<br>EBV reactivation is linked to immune suppression and may increase the risk of lymphoproliferative diseases and B-cell malignancies.<br>Prolonged EBV activity could lead to Hodgkin's lymphoma, peripheral B and T-cell lymphomas, and nasopharyngeal carcinoma | Rashes may occur, especially after taking ampicillin or amoxicillin.<br>Fatigue can persist for up to a year in some patients.<br>Complications may include hemolytic anemia, splenic rupture, hemophagocytic lymphohistiocytosis, and neurological problems.<br>EBV is associated with Burkitt's lymphoma.<br>EBV is linked to nasopharyngeal carcinoma.<br>EBV is connected to Hodgkin's lymphoma.<br>EBV can cause B-cell and T-cell lymphomas.<br>EBV is linked to oral hairy leukoplakia in immunocompromised individuals.<br>EBV can cause interstitial lymphocytic pneumonia.<br>EBV can lead to post-transplant lymphoproliferative disease.<br>EBV may contribute to mesenchymal tumors such as leiomyosarcoma. | COVID-19 may reactivate latent EBV, especially in severe cases.<br>Shared symptoms include fatigue, brain fog, headaches, confusion, and myalgia.<br>Both viruses disrupt mitochondrial and immune pathways, contributing to long COVID symptoms.<br>EBV reactivation occurred in 4.5% of COVID-19 patients. EBV-positive patients had higher fever incidence, lower albumin, lower platelet count, and longer hospital stay.<br>Long COVID and EBV share symptoms like fatigue, cognitive dysfunction, and sleep disturbances.<br>EBV reactivation is proposed as a common mechanism, potentially inducing acquired immunodeficiency.<br>Both conditions show immune dysregulation, chronic inflammation, and viral reactivation, suggesting similar underlying pathophysiology.<br>COVID-19 patients showed a 2-fold higher EBV reactivation rate (27.1%) than non-COVID patients (12.5%).<br>EBV reactivation may contribute to COVID symptoms, severity, and long |
|  |  |  |  | COVID, and was more common during the Omicron wave.<br>Shared symptoms and immune stress responses suggest possible overlap in post-viral complications.<br>The clinical presentation of these ulcers was similar to those caused by EBV, suggesting COVID-19 may trigger comparable immune responses or pathophysiological mechanisms.<br>A patient with mild COVID-19 developed worsening liver function, and tests confirmed EBV reactivation, which likely contributed to acute hepatitis.<br>EBV reactivation is frequently observed in long COVID and shares symptoms like fatigue, brain fog, sleep issues, myalgia, and GI upset.<br>EBV reactivation occurred in 65% of critically ill COVID-19 patients.<br>EBV/COVID-19 co-infection was associated with fever, higher CRP, and elevated AST, suggesting increased inflammation and potential disease severity.<br>EBV reactivation may serve as a marker of COVID-19 severity, possibly contributing to long COVID<br>symptoms, inflammation, and immune suppression.<br>EBV reactivation following SARS-CoV-2 infection is linked to increased inflammation, fatigue, and PASC (long COVID).<br>Both EBV and COVID-19 trigger NLRP3 inflammasome activation, contributing to persistent inflammation.<br>Those with EBV had higher rates of respiratory failure, ARDS, hypoproteinaemia, and elevated CRP and D-dimer, along with lower lymphocytes and albumin.<br>Shared symptoms included fatigue, myalgia, cognitive issues, fever, and pulmonary dysfunction.<br>EBV may contribute to long COVID via immune suppression, inflammation, and mitochondrial dysfunction, overlapping with autoimmune-like disorders.<br>Shared features include immune suppression, anemia, and hyperinflammation.<br>Both viruses showed symptom overlap including asthenia, sore throat, rash, anosmia, and ageusia. |
|  |  |  |  | Coinfection may cause diagnostic confusion and possibly aggravate clinical condition.<br>EBV reactivation is suggested as a potential contributor to long COVID symptoms like fatigue and neurocognitive dysfunction.<br>EBV and COVID-19 both increase the risk of periodontitis, possibly through immune modulation and inflammation.<br>EBV is associated with chronic and aggressive periodontitis, while COVID-19 may worsen periodontal disease and systemic outcomes via cytokine storms.<br>Suggested shared mechanisms include immune suppression and oral tissue vulnerability.<br>SARS-CoV-2 co-infection may trigger EBV lytic reactivation, leading to increased inflammation, fever, and immune suppression.<br>EBV reactivation has been linked to long COVID, and shared features include cytokine storms, CD8+ T cell loss, mitochondrial dysfunction, and upregulated ACE2 expression via EBV lytic genes. |
|  |  |  |  | SARS-CoV-2 and EBV share pathogenic traits, including molecular mimicry, autoimmunity induction, and inflammation.<br>Both viruses are linked to multiple sclerosis, chronic fatigue, and post-COVID syndromes.<br>Authors suggest COVID-19 may trigger EBV reactivation, contributing to gastrointestinal inflammation, long COVID symptoms, and slow recovery.<br>Both viruses linked to immune dysregulation and chronic inflammatory responses.<br>Post-COVID and post-EBV share persistent fatigue as a primary symptom. |
| 8 | Pseudomonas aeruginosa | Increased biofilm production in microgravity, making it more resistant to treatments.<br>Causes opportunistic infections, especially in burn victims and immunocompromised individuals<br>P. aeruginosa demonstrated differential regulation of 167 genes and 28 proteins under spaceflight conditions | Causes infection and bacteremia mainly in immunocompromised individuals.<br>Commonly infects the lower respiratory tract, leading to pneumonia.<br>Can cause necrotizing bronchopneumonia in cystic fibrosis patients.<br>Frequently responsible for ventilator-associated pneumonia in hospitals. | Review not conducted due to time constraints. |
|  |  | Increased production of virulence factors such as lecA, lecB, and rhlA, which are involved in bacterial adhesion, biofilm formation, and cytotoxicity<br>Spaceflight samples showed enhanced resistance to oxidative stress and biofilm formation, increasing the risk of persistent infections<br>Respiratory infections risk: *P. aeruginosa* has been found in astronaut microbiomes and is linked to lung infections in immunocompromised individuals<br>Impaired immune response under microgravity conditions led to increased bacterial load in blood.<br>Higher morbidity rates in irradiated and suspended mice due to reduced clearance of the infection | May lead to endocarditis, osteomyelitis, urinary tract infections, gastrointestinal infections, and meningitis.<br>Can cause septicemia (blood poisoning).<br>Often causes corneal infections in contact lens wearers, leading to scarring or vision loss.<br>Infects burn wounds, potentially causing abscesses, sepsis, and tissue discoloration.<br>Associated with swimmer's ear (otitis externa).<br>Has a mortality rate of around 30%, which may be higher depending on health conditions. |  |
| 9 | *Escherichia coli* | Increased resistance to antibiotics.<br>Altered gene expression in microgravity, leading to enhanced survival and potential virulence<br>Increased antibiotic resistance observed in *E. coli* isolates from astronauts. | Causes abrupt onset of watery diarrhea without blood, pus, or mucus.<br>Diarrhea is usually mild to moderate but can lead to severe fluid loss.<br>Low-grade fever, nausea, and abdominal pain may occur.<br>Severe dehydration may develop, especially in neonates and children. | Review not conducted due to time constraints. |
|  |  | Spaceflight conditions led to changes in bacterial phenotype, increasing its resistance to colistin and kanamycin.<br>Potential for opportunistic infections due to changes in intestinal flora | Illness typically lasts 2–5 days but can extend to 7 days or longer.<br>In severe cases, the disease can be life-threatening without treatment.<br>Estimated 800,000 deaths occur annually due to ETEC.<br>Most common in young children in developing countries and travelers from non-endemic regions. |  |
| 10 | Staphylococcus aureus | S. aureus was detected in the upper respiratory tract of astronauts.<br>In-flight cross-contamination between astronauts was observed.<br>Increase in opportunistic infections due to changes in bacterial flora caused by spaceflight<br>Staphylococcus aureus was not detected in the astronaut's nasal microbiome before flight but was present after 3 months aboard the ISS, suggesting possible microbial colonization from the spacecraft environment<br>Increased relative abundance of Staphylococcus aureus in the nasal microbiome could contribute to upper respiratory symptoms such as congestion, rhinitis, and sneezing, | Causes food poisoning with symptoms like nausea, vomiting, abdominal pain, and diarrhea.<br>Leads to skin infections such as pimples, impetigo, folliculitis, blisters, and abscesses.<br>Can cause cellulitis, erythema, tenderness, and mild fever from animal bites.<br>Responsible for scalded skin syndrome, especially in neonates and children.<br>May lead to necrotizing fasciitis, a rare but life-threatening condition.<br>Produces toxins that can cause toxic shock syndrome (TSS), a severe illness with high fever, rash, hypotension, and multi-organ failure. | Review not conducted due to time constraints. |
|  |  | which were commonly reported by astronauts | Can cause deep infections including endocarditis, peritonitis, necrotizing pneumonia, bacteremia, meningitis, osteomyelitis, and septic arthritis. |  |
| 11 | Klebsiella pneumoniae infections | Higher bacterial persistence in lung tissue under spaceflight-like conditions.<br>Increased pulmonary infection risk due to reduced granulocyte response | Causes nosocomial pneumonia, accounting for 7–14% of hospital-acquired cases.<br>Can lead to septicemia, urinary tract infections, and wound infections.<br>Commonly infects ICU patients and causes neonatal septicemias.<br>Responsible for community-acquired and hospital-acquired pneumonia with symptoms like fever, chills, and red currant jelly-like sputum.<br>May cause lung abscesses and necrosis of lung lobes.<br>Can lead to community-acquired meningitis and brain abscesses.<br>May cause pyogenic liver abscesses with symptoms like right upper quadrant pain, fever, and gastrointestinal discomfort.<br>Frequently causes urinary tract infections, including in patients with catheters. | Review not conducted due to time constraints. |
|  |  |  | Can result in other infections such as endocarditis, peritonitis, necrotizing fasciitis, and septic arthritis.<br>Klebsiella granulomatis causes donovanosis, a chronic ulcerative disease of the genitalia.<br>Infections are more common in immunocompromised individuals, especially those with diabetes, alcoholism, or chronic illness.<br>Infections can be severe and associated with high morbidity and mortality. |  |
| 12 | Staphylococcus epidermidis | Staphylococcus epidermidis was present throughout the mission but showed a shift in community composition | Pathogen Safety Data Sheet Not Available | Review not conducted due to time constraints. |

**Table 3:** Comparison of guidelines for managing airborne infectious diseases in space and on Earth, including COVID-19. This table summarizes key recommendations from studies on airborne pathogens in space, standard Earth-based biosafety measures, and existing COVID-19 guidelines. It includes containment protocols, infection prevention strategies, immune system monitoring, microbial surveillance, and therapeutic considerations. Guidelines were also aligned using the Canadian Biosafety Handbook based on Containment Level 2 and Risk Group 2 classifications.

| Guidelines for Airborne Infectious Diseases in Space | Guidelines for Airborne Infectious Diseases on Earth | Guidelines for COVID-19 on Earth |
| --- | --- | --- |
| • Emphasis on the need for in-depth understanding of infection mechanisms in space<br>• Potential use of nutritional countermeasures and probiotics for infection prevention during spaceflight.<br>• Highlighting the need to monitor opportunistic pathogens in space environments, as increased virulence may pose risks to immunocompromised astronauts. Further research recommended.<br>• Emphasizes NASA’s Health Stabilization Program (HSP) for infection prevention.<br>• Suggests vaccination protocols before space travel.<br>• Recommends in-flight isolation rooms and monitoring microbial changes during space missions.<br>• Recommends monitoring opportunistic pathogens in space environments.<br>• Suggests further research on microbial adaptability and infection risks for astronauts in microgravity conditions.<br>• Emphasizes the need for real-time microbial monitoring during space missions.<br>• Recommends biosensors for pathogen detection and antimicrobial resistance testing.<br>• Suggests evaluating astronaut immune function to mitigate infection risks in space.<br>• Recommends regular screening for microbial contamination before space travel.<br>• Suggests probiotic intake to maintain a healthy microbiome.<br>• Emphasizes the need for alternative therapies such as bacteriophage therapy and photodynamic therapy due to antibiotic resistance concerns in space.<br>• Suggests regular monitoring of *P. aeruginosa* in spacecraft water systems.<br>• Recommends biofilm disruption strategies to reduce infection risk in closed environments. | Based on All available Pathogen Safety Data Sheets, the pathogens studied in space are all classified as Containment Level 2 and Risk Group 2.<br>The Standard guidelines for these diseases according to Canadian Biosafety handbook are:<br>• Administrative controls (e.g., biosafety program management, training)<br>• Standardized procedures (e.g., work practices, personal protective equipment [PPE] use, and decontamination) that mitigate the risks associated with the activities conducted within the zone.<br>• Physical containment features include facility design (e.g., location of laboratory, surface finishes, access control) | Based on available Pathogen Safety Data Sheet on COVID-19, the pathogens studied in space are all classified as Containment Level 2 and Risk Group 2.<br>The Standard guidelines for these diseases according to Canadian Biosafety handbook are:<br>• Administrative controls (e.g., biosafety program management, training)<br>• Standardized procedures (e.g., work practices, personal protective equipment [PPE] use, and decontamination) that mitigate the risks associated with the activities conducted within the zone.<br>• Physical containment features include facility design (e.g., location of laboratory, |
| • Highlights the need for better antimicrobial resistance testing in space<br>• Recommends monitoring astronauts for viral reactivation pre- and post-flight.<br>• Suggests antiviral prophylaxis and immune-boosting interventions to prevent EBV-related complications in space.<br>• Calls for further research into radiation-induced viral reactivation mechanisms<br>• Monitoring and controlling airborne pathogens in spacecraft water and air systems.<br>• Developing countermeasures to prevent bacterial persistence in space habitats.<br>• Studying immune system responses further to mitigate infection risks during extended missions<br>• Monitoring for VZV reactivation in astronauts before, during, and after space missions.<br>• Suggests antiviral prophylaxis for astronauts with a history of VZV infection.<br>• Calls for further research into the long-term effects of VZV reactivation in space conditions<br>• Suggests continuous monitoring of Salmonella on spacecraft surfaces.<br>• Recommends studying astronaut microbiome shifts to assess infection risks in space.<br>• Calls for targeted countermeasures to reduce spaceflight-enhanced virulence<br>• Suggests continuous monitoring of Salmonella virulence factors in space environments.<br>• Recommends developing infection control strategies for long-duration missions.<br>• Calls for targeted antimicrobial measures to counteract spaceflight-induced bacterial virulence<br>• Continuous microbial monitoring of astronaut microbiomes before, during, and after spaceflight. | • Provision of biosafety equipment, such as primary containment devices (e.g., biological safety cabinets [BSCs]) for certain activities. | surface finishes, access control)<br>• Provision of biosafety equipment, such as primary containment devices (e.g., biological safety cabinets [BSCs]) for certain activities. |
| • Suggests further research on microbial transmission risks in space habitats.<br>• Calls for improved environmental controls to limit bacterial colonization in spacecraft |  |  |

To better conceptualize and apply these guidelines in a space mission context, the procedures were categorized into four main operational areas: pre-flight measures, in-flight prevention strategies, astronaut health and immune system support, and in-flight isolation and emergency response protocols (Table 4). This framework will help in organizing the response to infectious disease threats in a way that addresses both prevention and mitigation, while also supporting astronaut health throughout the mission duration.

**Table 4:** Recommended COVID-19 prevention and response strategies for space missions. The table outlines comprehensive measures including pre-flight vaccination and screening, in-flight environmental and health monitoring, immune system support, and isolation protocols.

|  |  |
| --- | --- |
| **Pre-Flight Measures:** | **Vaccination:** Ensure astronauts are vaccinated against COVID-19 before traveling to space.<br>**Screening:** Astronauts should be tested for COVID-19 before the mission. This includes regular checks for symptoms and testing as needed.<br>**Health Monitoring:** Continuous health checks, including temperature and symptom screening, should be done before and during the mission. |
| **In-Flight Prevention:** | **Air and Water Systems:** Ensure spacecraft have proper filtration systems to prevent airborne pathogens and maintain clean water systems.<br>**Medical Support:** Telemedicine should be available to consult with Earth-based doctors if an astronaut shows symptoms of COVID-19.<br>**Decontamination:** Routine cleaning and disinfection of surfaces in the spacecraft and equipment should be conducted using appropriate procedures and PPE. |
| **Astronaut Health and Immune Support:** | **Immune Function Monitoring:** Astronauts' immune health should be regularly monitored to detect any changes that might increase infection risk.<br>**Nutritional Support:** Provide astronauts with dietary supplements, including probiotics, to support immune function and prevent infections.<br>**Mental Health:** Ensure astronauts have mental health support to manage stress, which can affect the immune system. |
| **In-Flight Isolation and Response:** | **Isolation Areas:** If an astronaut shows signs of infection, isolate them in a designated area within the spacecraft.<br>**Environmental Controls:** Regular microbial monitoring of the spacecraft environment is essential, especially for airborne viruses like SARS-CoV-2.<br>**Emergency Response:** Prepare for contingencies such as extended isolation or quarantine if an outbreak occurs. |

## Discussion

Since the first human ventured into space in 1961, space travel has evolved from a daring exploration to a burgeoning industry, with 643 individuals having traveled beyond Earth’s atmosphere to date. Some of these astronauts have even completed as many as seven missions to space over a short span, highlighting the increasing frequency of space travel [19]. This growing interest has spurred investments from companies like SpaceX, Virgin Galactic, and Blue Origin, which are collectively driving advancements in space exploration and research [43]. In addition, countries such as India and China have multiple future space missions planned; with China planning space tourism and India planning to launch the world’s third space station [2, 21]. Despite these technological and commercial advancements, the health risk and long-term implications of infectious diseases during space missions remains a pressing concern. The Apollo 7 mission, for example, experienced a significant setback when all three astronauts contracted the respiratory viral infection, which rapidly spread in the confined spacecraft and disrupted their ability to work effectively as a team [36]. Such events emphasize the critical need to address and mitigate the potential risks of highly contagious diseases, such as COVID-19, in the unique microgravity environment involved in space travel. The rise of the COVID-19 pandemic began in March 2020, with the spread of the disease to all parts of the world. COVID-19 infected over 776 million people worldwide, resulting in the deaths of over 1.8 million people [52]. The disease is caused by a viral vector, SARS-CoV-2, which is a single-stranded RNA virus [8]. There are multiple ways COVID-19 can be transmitted. However, it is primarily transmitted through respiratory droplets and aerosols released when an infected person coughs, sneezes, or speaks [8]. With the uniqueness of microgravity environment, microdroplets even after sneezing will stay in the artificial environment, thus making it easy for another astronaut to breaths it in. Conducting a study to understand the health risks posed by COVID-19 in space is essential for ensuring mission success and preventing unnecessary fatalities during long-duration space travel. This study aims to address the lack of research on COVID-19 in space by examining the behavior of other airborne infectious diseases in microgravity and comparing their effects on Earth and in space. The objectives include conducting systematic reviews to identify and analyze studies on airborne diseases, comparing the effects of these diseases with COVID-19, and predicting the potential risks of COVID-19 in space. The goal is to develop an evidence-based preparedness plan by integrating space and Earth guidelines for managing airborne infections, reducing the risk of COVID-19 transmission during space missions.

With the potential emergence of a $1.8 trillion market, companies like SpaceX, Virgin Galactic, and Blue Origin are driving significant interest and investment in space exploration [23, 43]. However, this increase in space travel also introduces greater liability for corporations. For instance, in 2005, there were 7,472 trials held against businesses, 1,863 of which were related to premises liability [25]. Failing to analyze and address the risks of COVID-19 could lead to similar liabilities for companies operating in the space travel sector. Furthermore, understanding these risks is vital for the success of future missions. With only seven professionals aboard spacecraft at a time and limited resources available in space, preventing the spread of COVID-19 becomes critical. This resource scarcity is further exacerbated by the constraints of space and mass aboard spacecraft [18]. Our findings could lead to innovations in spacecraft design, astronaut training, and disease prevention protocols, ultimately benefiting commercial space enterprises and international space agencies. By systematically assessing current literature, we can contribute to the development or enhancement of space travel guidelines, enabling governments and corporations to better prepare for and ensure safe and successful missions.

The study was conducted over a period of four months and utilized both publicly accessible databases and student access provided by the Kwantlen Polytechnic University Library to retrieve articles behind paywalls [24]. Through systematic reviews, nine diseases were identified for analysis. Of the nine diseases reviewed, Salmonella typhimurium, Serratia marcescens, and Epstein-Barr virus (EBV) were selected for comparison. Salmonella typhimurium and Serratia marcescens were primarily chosen due to the limited number of available studies, whereas EBV was selected for its viral nature. Additionally, COVID-19 patients exhibit an increased incidence of EBV reactivation, further strengthening the association between the two viruses [3]. From the extraction and comparison, six tentative effects of COVID-19 in space were identified. Two of these Immune Suppression & Reactivation and Neurocognitive & Systemic Symptoms were derived from the comparison between EBV and COVID-19. EBV reactivation has been observed in space, where it suppresses immune function and affects multiple organ systems [31]. The prevalence of post-COVID conditions, including long COVID where signs and symptoms persist or reappear after four or more weeks further strengthens this connection [16]. One effect, Increased Virulence & Growth Potential, was identified based on the relationship between COVID-19 and Serratia marcescens, supported by findings that demonstrate increased virulence of Serratia marcescens under spaceflight conditions [15]. The remaining three effects Inflammatory Response, Gastrointestinal & Pulmonary Risk, and Greater Severity & Mortality Risk were general observations drawn from both EBV and Serratia marcescens. Both EBV and COVID-19 are known to trigger inflammasome activation, contributing to persistent inflammation on Earth [42]. Given that EBV exhibits similar inflammatory responses in space and increases gastrointestinal and pulmonary risks, it is reasonable to predict that COVID-19 may behave in a similar manner in microgravity environments [31].

While the guidelines were initially collected generically for all airborne infectious diseases in space, it was later revealed during the study that all the pathogens identified in the initial phase belong to Containment Level 2, Risk Group 2, according to the Pathogen Safety Data Sheet. COVID-19 also falls under the same Containment Level 2 and Risk Group 2 classification [41]. Based on this, guidelines were modeled to prevent the spread of COVID-19 in space by combining the generic recommendations for Containment Level 2 and Risk Group 2 from the Canadian Biosafety Handbook with the guidelines previously collected from studies on airborne infectious diseases in space. When comparing this approach to the NASA Space Flight Human-System Standard, it was found that most of the suggested guidelines are already in place. This further validates the viability of the method used in this study for guideline development and indicates that it is aligned with current best practices.

This research allows for the inference of potential behaviors that COVID-19 may exhibit in space or microgravity. Such insights may contribute to making future space travel safer by helping space agencies and astronauts better understand the associated risks. The identification of tentative health effects, such as immune suppression and increased virulence, underscores the importance of incorporating targeted prevention and response strategies into spaceflight protocols. These findings can inform the refinement of medical protocols across all phases of a mission: pre-flight, in-flight, and post-flight by guiding screening, isolation, and immune monitoring procedures specific to COVID-19 and similar airborne pathogens. As commercial space tourism expands, this knowledge also becomes critical for developing risk management strategies for non-career astronauts who may have varying levels of medical fitness and exposure history.

Due to time limitations, ScienceDirect was not used during the initial collection of studies on airborne infectious diseases. However, it is a significant source of research and is expected to greatly expand the scope of future investigations. Similarly, the comparison process was limited to Salmonella typhimurium, Serratia marcescens, and Epstein-Barr virus (EBV) due to time constraints. Although Varicella-Zoster Virus (VZV) was also shortlisted, given its viral nature, it could not be included in the analysis for the same reason. Notably, VZV reactivation has been reported following COVID-19 vaccination [29], suggesting a potential area for future exploration. Additionally, since only PubMed was used for this phase of the study, incorporating research from other databases would further improve both the specificity and comprehensiveness of the findings.

The approach utilized in this paper will also enable future studies to compare and estimate the risks of disease in space without relying on direct samples or waiting for an outbreak. The technique involves utilizing details of diseases studied in extreme environments, comparing their effects in regular environments, and then applying a comparative approach to the disease of interest. This allows for more accurate risk analysis and understanding of health effects in extreme environments. This methodological advancement could be pivotal for analyzing health risks in other extreme or inaccessible settings. Additionally, future research could expand on this foundation to examine the effects of long COVID-19 in space, offering insights into the potential complications of this condition under microgravity. Long COVID-19 refers to symptoms persisting for more than 12 weeks after the onset of infection [39]. By addressing these knowledge gaps, this research will contribute to a broader understanding of infectious disease behavior in unique environments and help pave the way for safer and more sustainable human presence beyond Earth.

## Acknowledgement

First and foremost, I would like to express my sincere gratitude to Kwantlen Polytechnic University, particularly the Faculty of Science and the Department of Biology and Health Science, for providing the academic environment and resources necessary to complete this research. I am especially thankful to my project supervisor, Dr. Barnabe D. Assogba, for his invaluable guidance, encouragement, and support throughout the course of this study. I would also like to thank my Honours course instructor, Dr. Layne Myhre, for his continuous support in developing my skills in literature review and scientific writing. A special thanks to Eddie Han for his assistance in the assessment of articles, and to all my peers for their constructive feedback and encouragement during the research process. This research was supported by the Student Research & Innovation Grant (SRIG) #104486 (Stream 2) from Kwantlen Polytechnic University (KPU).

## References

1. Araújo, M. B.; Naimi, B. Spread of SARS-Cov-2 Coronavirus Likely Constrained by Climate. medRxiv 2020, DOI: 10.1101/2020.03.12.20034728.

2. Bansal, R. India Will Soon Be the Third Country with Its Own Space Station: A Historic LEAP in Space Exploration. YourStory, September 24, 2024. Retrieved November 19, 2024, from Link.

3. Bernal, K. D.; Whitehurst, C. B. Incidence of Epstein-Barr Virus Reactivation Is Elevated in COVID-19 Patients. Virus Res. 2023, 334, 199157. DOI: 10.1016/j.virusres.2023.199157.

4. Brosnahan, S. B.; Jonkman, A. H.; Kugler, M. C.; Munger, J. S.; Kaufman, D. A. COVID-19 and Respiratory System Disorders. Arterioscler. Thromb. Vasc. Biol. 2020, 40(11), 2586–2597. DOI: 10.1161/atvbaha.120.314515.

5. Brown, E. Boosting Rocket Reliability at the Material Level. MIT News, November 28, 2023. Retrieved December 9, 2024, from Link.

6. Cannell, J. J.; Vieth, R.; Umhau, J. C.; Holick, M. F.; Grant, W. B.; Madronich, S.; Garland, C. F.; Giovannucci, E. Epidemic Influenza and Vitamin D. Epidemiol. Infect. 2006, 134(6), 1129–1140. DOI: 10.1017/s0950268806007175.

7. CDC. Signs and Symptoms of Long COVID. Centers for Disease Control and Prevention, September 17, 2024. Retrieved December 8, 2024, from [Link](https://www.cdc.gov/covid/long-term-effects/long-covid-signs-symptoms.html?form=MG0AV3).

8. Chakraborty, I.; Maity, P. COVID-19 Outbreak: Migration, Effects on Society, Global Environment and Prevention. Sci. Total Environ. 2020, 728, 138882. DOI: 10.1016/j.scitotenv.2020.138882.

9. Chen, B.; Liang, H.; Yuan, X.; Hu, Y.; Xu, M.; Zhao, Y.; Zhang, B.; Tian, F.; Zhu, X. Roles of Meteorological Conditions in COVID-19 Transmission on a Worldwide Scale. medRxiv 2020, DOI: 10.1101/2020.03.16.20037168.

10. CSIS Aerospace Security Project. Cumulative Number of Human Visits to Space. Our World in Data, 2022. Retrieved October 13, 2024, from [Link](https://ourworldindata.org/grapher/cumulative-space-visits?country=OWID_WRL%7EJPN%7ERUS%7ECAN%7EUSA).

11. CSIS Aerospace Security Project. Human Visits to Space per Year. Our World in Data, 2022. Retrieved October 13, 2024, from [Link](https://ourworldindata.org/grapher/annual-space-visits).

12. Dahl, C.; Gilbert, B.; Lange, I. Mineral Scarcity on Earth: Are Asteroids the Answer? Miner. Econ. 2020, 33(1–2), 29–41. DOI: 10.1007/s13563-020-00231-6.

13. Diaz, J. V.; Soriano, J. B. A Delphi Consensus to Advance on a Clinical Case Definition for Post COVID-19 Condition: A WHO Protocol. Lancet Infect. Dis. 2021, 22(4). DOI: 10.21203/rs.3.pex-1480/v1.

14. Edalatifard, M.; Rahimi, B.; Vesal, A. Coronavirus and Its Effect on the Respiratory System: Is There Any Association between Pneumonia and Immune Cells. J. Family Med. Prim. Care 2020, 9(9), 4729. DOI: 10.4103/jfmpc.jfmpc_763_20.

15. Gilbert, R.; Torres, M.; Clemens, R.;, et al. Spaceflight and Simulated Microgravity Conditions Increase Virulence of Serratia marcescens in the Drosophila melanogaster Infection Model. npj Microgravity 2020, 6(1). DOI: 10.1038/s41526-019-0091-2.

16. Greenhalgh, T.; Sivan, M.; Perlowski, A.; Nikolich, J. Z. Long COVID: A Clinical Update. Lancet 2024, 404(10453).

17. Guo, Z.; Tang, Y.; Zhang, Z.;, et al. COVID-19: From Immune Response to Clinical Intervention. *Precis*. Clin. Med. 2024, 7(3). DOI: 10.1093/pcmedi/pbae015.

18. Hepp, A. F.; Palaszewski, B. A.; Colozza, A. J.; Landis, G. A.; Jaworske, D. A.; Kulis, M. J. In-Situ Resource Utilization for Space Exploration: Resource Processing, Mission-Enabling Technologies, and Lessons for Sustainability on Earth and Beyond. 12th International Energy Conversion Engineering Conference, 2014. DOI: 10.2514/6.2014-3761.

19. Hobbs, Z. How Many People Have Gone to Space? *Astronomy Magazine*, November 17, 2023. Retrieved November 19, 2024, from Link.

20. 20. Hurley, S. Where Does Space Begin? Explaining Science, January 8, 2019. Retrieved April 19, 2025, from [Link](https://explainingscience.org/2019/01/01/where-does-space-begin/).

21. Jones, A. China Seeks Its Own Apollo Moment – and More. SpaceNews, June 5, 2024. Retrieved November 19, 2024, from Link.

22. Jones, C. W.; Overbey, E. G.; Lacombe, J.;, et al. Molecular and Physiological Changes in the SpaceX Inspiration4 Civilian Crew. Nature 2024, 632(8027), 1155–1164. DOI: 10.1038/s41586-024-07648-x.

23. 23. Khlystov, N.; Markovitz, G. Space Is Booming. Here’s How to Embrace the $1.8 Trillion Opportunity. World Economic Forum, April 8, 2024. Retrieved December 9, 2024, from [Link](https://www.weforum.org/stories/2024/04/space-economy-technology-invest-rocket-opportunity/).

24. Kwantlen Polytechnic University. KPU Library. Retrieved from [Link](https://www.kpu.ca/library).

25. Langton, L.; Cohen, T. H. Civil Bench and Jury Trials in State Courts, 2005. *U.S. Department of Justice*, 2009. Retrieved from [Link](https://bjs.ojp.gov/content/pub/pdf/cbjtsc05.pdf?form=MG0AV3).

26. Li, J.; Huang, D. Q.; Zou, B.;, et al. Epidemiology of COVID-19: A Systematic Review and Meta-Analysis of Clinical Characteristics, Risk Factors, and Outcomes. J. Med. Virol. 2020, 93(3), 1449–1458. DOI: 10.1002/jmv.26424.

27. Liu, Q.; Zhou, R.; Zhao, X.; Chen, X.; Chen, S. Acclimation During Space Flight: Effects on Human Emotion. Mil. Med. Res. 2016, 3(1). DOI: 10.1186/s40779-016-0084-3.

28. Lowen, A. C.; Mubareka, S.; Steel, J.; Palese, P. Influenza Virus Transmission Is Dependent on Relative Humidity and Temperature. PLoS Pathog. 2007, 3(10), e151. DOI: 10.1371/journal.ppat.0030151.

29. Martínez-Reviejo, R.; Tejada, S.; Adebanjo, G. A.;, et al. Varicella-Zoster Virus Reactivation Following Severe Acute Respiratory Syndrome Coronavirus 2 Vaccination or Infection: New Insights. Eur. J. Intern. Med. 2022, 104, 73–79. DOI: 10.1016/j.ejim.2022.07.022.

30. Mecenas, P.; Bastos, R. T.; Vallinoto, A. C.; Normando, D. Effects of Temperature and Humidity on the Spread of COVID-19: A Systematic Review. PLoS One 2020, 15(9), e0238339. DOI: 10.1371/journal.pone.0238339.

31. Mehta, S.; Bloom, D.; Plante, I.;, et al. Reactivation of Latent Epstein-Barr Virus: A Comparison after Exposure to Gamma, Proton, Carbon, and Iron Radiation. Int. J. Mol. Sci. 2018, 19(10), 2961. DOI: 10.3390/ijms19102961.

32. Montesinos, C. A.; Khalid, R.; Cristea, O.;, et al. Space Radiation Protection Countermeasures in Microgravity and Planetary Exploration. Life 2021, 11(8), 829. DOI: 10.3390/life11080829.

33. Mudgal, S. K.; Gaur, R.; Rulaniya, S.;, et al. Pooled Prevalence of Long COVID-19 Symptoms at 12 Months and Above Follow-Up Period: A Systematic Review and Meta-Analysis. Cureus 2023. DOI: 10.7759/cureus.36325.

34. 34. Ostovar, M. About Apollo 7, the First Crewed Apollo Space Mission. NASA, October 10, 2023. Retrieved December 10, 2024, from [Link](https://www.nasa.gov/missions/apollo/about-apollo-7-the-first-crewed-apollo-space-mission/).

35. Overbey, E. G.; Kim, J.; Tierney, B. T.;, et al. The Space Omics and Medical Atlas (SOMA) and International Astronaut Biobank. Nature 2024, 632(8027), 1145–1154. DOI: 10.1038/s41586-024-07639-y.

36. Pavlečić, B.; Runzheimer, K.; Siems, K.;, et al. Spaceflight Virology: What Do We Know About Viral Threats in the Spaceflight Environment? Astrobiology 2022, 22(2), 210–224. DOI: 10.1089/ast.2021.0009.

37. Prisk, G. K. Pulmonary Challenges of Prolonged Journeys to Space: Taking Your Lungs to the Moon. Med. J. Aust. 2019, 211(6), 271–276. DOI: 10.5694/mja2.50312.

38. Public Health Agency of Canada. COVID-19: Symptoms, Treatment, What to Do If You Feel Sick. *Canada.ca*, January 27, 2023. Retrieved December 8, 2024, from [Link](https://www.canada.ca/en/public-health/services/diseases/2019-novel-coronavirus-infection/symptoms.html).

39. Public Health Agency of Canada. Post-COVID-19 Condition (Long COVID). *Canada.ca*, October 22, 2024. Retrieved December 10, 2024, from [Link](https://www.canada.ca/en/public-health/services/diseases/2019-novel-coronavirus-infection/symptoms/post-covid-19-condition.html).

40. Public Health Agency of Canada. Pathogen Safety Data Sheets. *Canada.ca*, February 5, 2025. Retrieved April 21, 2025, from [Link](https://www.canada.ca/en/public-health/services/laboratory-biosafety-biosecurity/pathogen-safety-data-sheets-risk-assessment.html).

41. Public Health Agency of Canada. Severe Acute Respiratory Syndrome Coronavirus-2 (SARS-CoV-2): Pathogen Safety Data. *Canada.ca*, April 13, 2023. Retrieved April 19, 2025, from [Link](https://www.canada.ca/en/public-health/services/laboratory-biosafety-biosecurity/pathogen-safety-data-sheets-risk-assessment/severe-acute-respiratory-syndrome-coronavirus-2.html).

42. Rousseau, B. A.; Bhaduri-McIntosh, S. Inflammation and Epstein–Barr Virus at the Crossroads of Multiple Sclerosis and Post-Acute Sequelae of COVID-19 Infection. Viruses 2023, 15(4), 949. DOI: 10.3390/v15040949.

43. Schweinsberg, S.; Fennell, D. Space Tourism: A Historical and Existential Perspective. Sustainability 2023, 16(1), 79. DOI: 10.3390/su16010079.

44. Sibonga, J. D.; Evans, H. J.; Smith, S. A.;, et al. Risk of Bone Fracture Due to Spaceflight-Induced Changes to Bone. NASA, 2017. Retrieved from [Link](https://humanresearchroadmap.nasa.gov/Evidence/reports/Fracture.pdf).

45. Singh, D.; Mathioudakis, A. G.; Higham, A. Chronic Obstructive Pulmonary Disease and COVID-19: Interrelationships. Curr. Opin. Pulm. Med. 2021, 28(2), 76–83. DOI: 10.1097/MCP.0000000000000834.

46. Smith, S. M.; Heer, M.; Shackelford, L. C.;, et al. Bone Metabolism and Renal Stone Risk During International Space Station Missions. Bone 2015, 81, 712–720. DOI: 10.1016/j.bone.2015.10.002.

47. Sutradhar, J.; Sarkar, B. R. The Effect on the Immune System in the Human Body Due to COVID-19: An Insight on Traditional to Modern Approach as a Preventive Measure. J. Pharmacopuncture 2021, 24(4), 165–172. DOI: 10.3831/kpi.2021.24.4.165.

48. Targa, A. D.; Benítez, I. D.; Moncusí-Moix, A.;, et al. Decrease in Sleep Quality During COVID-19 Outbreak. Sleep Breath. 2020, 25(2), 1055–1061. DOI: 10.1007/s11325-020-02202-1.

49. 49. Taveau, J. NASA Accelerates Space Exploration, Earth Science for All in 2024. *NASA*, December 6, 2024. Retrieved December 8, 2024, from [Link](https://www.nasa.gov/news-release/nasa-accelerates-space-exploration-earth-science-for-all-in-2024/).

50. Wang, J.; Tang, K.; Feng, K.;, et al. Impact of Temperature and Relative Humidity on the Transmission of COVID-19: A Modelling Study in China and the United States. BMJ Open 2021, 11(2), e043863. DOI: 10.1136/bmjopen-2020-043863.

51. WHO Data. WHO COVID-19 Dashboard. World Health Organization, January 12, 2020. Retrieved October 8, 2024, from [Link](https://data.who.int/dashboards/covid19/cases?n=c).

52. World Health Organization. The True Death Toll of COVID-19. World Health Organization, April 2022. Retrieved October 15, 2024, from [Link](https://www.who.int/data/stories/the-true-death-toll-of-covid-19-estimating-global-excess-mortality).

53. World Health Organization. Tracking SARS-CoV-2 Variants. World Health Organization, December 2, 2024. Retrieved December 8, 2024, from [Link](https://www.who.int/activities/tracking-SARS-CoV-2-variants/?form=MG0AV3).

54. World Health Organization. Naming the Coronavirus Disease (COVID-19) and the Virus That Causes It. World Health Organization, n.d. Retrieved October 8, 2024, from [Link](https://www.who.int/emergencies/diseases/novel-coronavirus-2019/technical-guidance/naming-the-coronavirus-disease-(covid-2019)-and-the-virus-that-causes-it).

